# Global Cropland Connectivity: A Risk Factor for Invasion and Saturation by Emerging Pathogens and Pests

**DOI:** 10.1101/106542

**Authors:** Y. Xing, J. F. Hernandez Nopsa, K. F. Andersen, J. Andrade-Piedra, F. D. Beed, G. Blomme, M. Carvajal-Yepes, D. L. Coyne, W. J. Cuellar, G. A. Forbes, J. F. Kreuze, J. Kroschel, P. L. Kumar, J. P. Legg, M. Parker, E. Schulte-Geldermann, K. Sharma, K. A. Garrett

## Abstract

The geographic pattern of cropland is an important risk factor for invasion and saturation by crop-specific pathogens and arthropods. Understanding cropland networks supports smart pest sampling and mitigation strategies. We evaluate global networks of cropland connectivity for key vegetatively-propagated crops (banana and plantain, cassava, potato, sweetpotato, and yam) important for food security in the tropics. For each crop, potential movement between geographic location pairs was evaluated using a gravity model, with associated uncertainty quantification. The highly-linked hub and bridge locations in cropland connectivity risk maps are likely priorities for surveillance and management, and for tracing intra-region movement of pathogens and pests. Important locations are identified beyond those locations that simply have high crop density. Cropland connectivity risk maps provide a new risk component for integration with other factors – such as climatic suitability, genetic resistance, and trade routes – to inform Pest Risk Assessment and mitigation.

---

Plant diseases and pests are major threats to food security and wildlands conservation (Aguayo et al. 2014, Anderson et al. 2004, Fisher et al. 2012, Gonthier and Garbelotto 2013, Woolhouse et al. 2005). Understanding which geographic areas have a high risk of pathogen and arthropod pest invasion is an important first step to designing sampling and mitigation strategies (Fears et al. 2014). Climate effects are one component of this risk, and are commonly addressed in species distribution models (Anderson et al. 2004, Bebber et al. 2013, Elith and Leathwick 2009, Garrett et al. 2006, Garrett et al. 2014, Hernandez Nopsa et al. 2014, Jeger and Pautasso 2008, Kroschel et al. 2016, Rodoni 2009, Rosenzweig et al. 2001). Another important risk component is the structure of trade routes, through which pathogens and pests may move (Anderson et al. 2004, Bebber et al. 2014, Nakato et al. 2013). Habitat connectivity represents a third component that, integrated with these other risk factors, and potentially other factors such as deployment of resistance, can provide a more complete invasion risk assessment. The connectivity of cropland regions helps to determine whether invasive species that are dependent on crops will become established before effective actions can be taken to mitigate them (Margosian et al. 2009, Sutherst 2014). Incorporating cropland connectivity risk with other risk factors for invasion supports a number of integrated pest management and pest risk assessment strategies, from improved methods for detecting and mitigating new invasives, to ongoing improvements in policy (Leung et al. 2015, Margosian et al. 2009, With 2004).

The invasion of species into new regions is a common research focus, but less attention is given to the process of saturation (Cornell and Lawton 1992, Fox et al. 2000, Lion and Gandon 2009). Bebber et al. (2014) considered saturation in terms of the fraction of potentially habitable regions that are already occupied by a pest species. Similarly, here we define saturation as the process by which a species “fills in” a region, to occupy more and more of the potential habitat within the region. Defining the difference between invasion and saturation is often a question of the spatial resolution and extent being considered (Fig. 1). From the standpoint of pathogen and arthropod management, a pest may have already invaded a region and be considered endemic, while at the same time there may be some fields it has never reached, and its population may frequently be suppressed by factors such as extreme weather conditions so that it must “re-saturate”. Some pathogens continue to “re-emerge” at different time points (Rugalema et al. 2009, Vurro et al. 2010), such as *Phytophthora infestans* (Fry et al. 2015). For pathogens like *P. infestans*, initial inoculum can be limiting. For example, unusually abundant initial inoculum probably played a key role in the devastating 2009-2010 late blight epidemic in tomato in the northeastern US (Fry et al. 2013). Similarly, high inoculum associated with synergistic virus species interactions was a key driving factor behind the rapid spread of the cassava mosaic disease pandemic in Africa in the 1990s (Harrison et al. 1997, Legg et al. 2006). High cropland connectivity is a risk factor for saturation and reemergence, as well as novel invasions, when pathogens and arthropods spread from refugia or nearby regions after limiting weather conditions.

**Figure 1.**
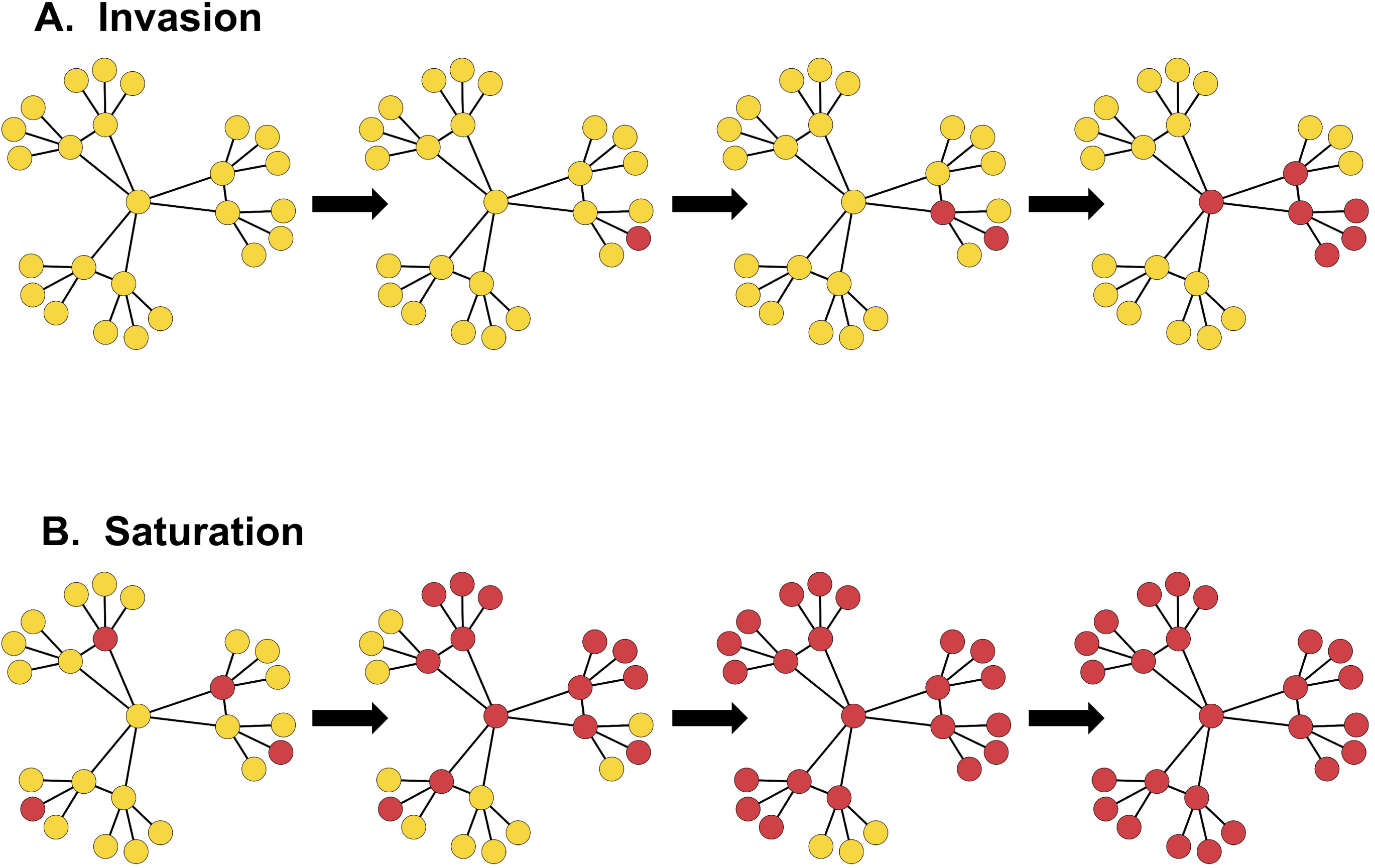
Contrasting the processes of invasion and saturation over time, where the pathogen or pest species is present in red nodes and absent in orange nodes. A. In an invasion, a region is initially free of the species. During the process of invasion, the species enters the region and spreads over time. B. In the process of saturation, a species is already present in a region but not in all potential locations. A restricted subset of nodes in the region may act as refugia for overwintering or oversummering, or for persistence of the species during years with weather less conducive to the species. From these locations, the species can spread to linked nodes when conditions are more conducive. Note that the difference between invasion and saturation is partially a difference in the spatial resolution being considered. For consideration of the risk due to cropland connectivity, even when a species is already present in the region, higher cropland connectivity increases the risk of saturation.

Network analysis offers a number of tools for understanding the strengths and vulnerabilities of network structures. In a geographic network analysis of species invasion or saturation, nodes represent geographic locations and the links between nodes represent functions such as the probability of movement of a pathogen or pest between the nodes. Characterizing the network structure of cropland areas acting as sinks or sources can inform the selection of key nodes for surveillance, mitigation, and management improvement. Nodes that are linked to many other nodes (nodes that have high degree) and nodes acting as bridges between cropland regions (nodes with high betweenness centrality) may be particularly important for the spread of pathogens and pests and are important for evaluating invasion risk (Hernandez Nopsa et al. 2015, Margosian et al. 2009). Network traits such as centrality (how important a particular node or link is (Newman 2010)), local cohesiveness (how well connected a subset of nodes is compared to their connection to other subsets of nodes (Kolaczyk 2009)), and affinity (degree of tendency for nodes to be linked with other nodes of similar centrality (Barrat et al. 2004)) can help to identify locations in networks that may be priorities for attention.

One problem for evaluating the effect of cropland connectivity, in general or for a particular pathogen or arthropod pest, is lack of information about current distribution and dispersal probabilities to parameterize dispersal risk models. More general risk evaluations can draw on models that have proven useful across multiple systems. The inverse power law function is commonly used to model pathogen dispersal. Parameter estimates for six cases – including plant and human pathogens and distances ranging from experimental field plots (32 m) to continental-scale (9329 km) – ranged from 1.75 to 2.36 (Mundt et al. 2009a). For cucurbit downy mildew, the observed maximum annual disease spread distance ranged from 1,914 km to 2,221 km across seven years, with inverse power law parameter estimates approximately 2 or more (Ojiambo et al. 2017). Gravity models are frequently used to describe the risk of movement between two locations, in applications including zoology, ecology, and epidemiology (Jongejans et al. 2014). In dispersal events, the risk of movement between two locations is often a function of the product of the amount of inoculum potentially produced at the source location and the amount of potential host material at the sink location, as in a gravity model (Jongejans et al. 2014, Sutrave et al. 2012, Xia et al. 2004).

Cassava mosaic disease (CMD) is an important example of the likely role of cropland connectivity in the spread of a plant disease epidemic (Fig. 2). Cassava mosaic begomoviruses (CMBs) cause CMD (Bock and Woods 1983), one of the most damaging constraints to cassava production in Sub-Saharan Africa. Losses have been estimated at more than US$1 billion per year (Legg et al. 2006). CMBs are dispersed via infected planting material and are vectored by the whitefly, *Bemisia tabaci* (Dubern 1994). Severe disease results when there is co-infection of cassava plants with African cassava mosaic virus (ACMV) and East African cassava mosaic virus (EACMV). A pandemic of severe CMD resulted from rapid spread of these synergistic mixed infections (Harrison et al. 1997, Legg et al. 2011). A heightened risk of disease spread was predicted for areas of widespread cassava cultivation, and slower spread was anticipated where there were major topographical barriers, such as lakes or dense forests (Legg 1999, Legg 2010). More specifically, contrasting rates of CMD spread down the east and west sides of Lake Victoria have been attributed to more contiguous cultivation of cassava on the western side of the Lake, and the physical barrier imposed by the Winam Gulf on the eastern side of Lake Victoria in western Kenya. The absence of cassava in central Kenya and central regions of Tanzania likely served as a barrier preventing the spread of the pandemic associated with EACMV-UG to coastal regions of East Africa (Szyniszewska et al. 2017).

**Figure 2.**
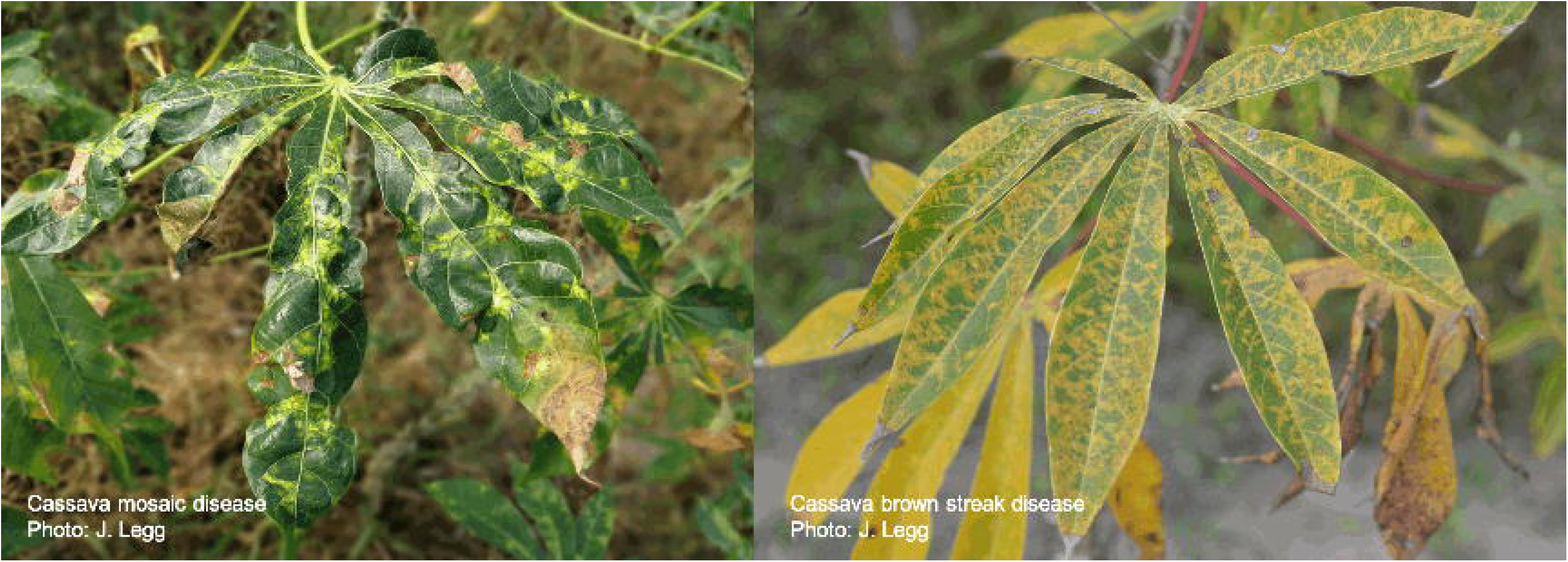
Cassava mosaic disease in Tai Ninh, Vietnam, and cassava brown streak disease in Mkuranga, Coast Province, Dar es Salaam.

We evaluate the global cropland connectivity risk associated with cassava and four other crops of particular importance to food security for smallholder farmers in the tropics. These crops are vegetatively-propagated, with the associated high risk of transmission of pests and diseases through planting materials. Cropland connectivity captures some elements of the risk of transmission through movement of pathogens and pests independent of crop plants (through flight or passive dispersal in wind, for example), as well as risk due to movement of planting materials and farm equipment. Because information about relevant dispersal kernels is often unavailable, uncertainty quantification (sensitivity analysis) may be needed to understand how dispersal parameters influence estimates. The objectives of this paper are to i) characterize the network structure of global cropland for banana and plantain, cassava, potato, sweetpotato, and yam (*Dioscorea* spp.), ii) evaluate the network structure in terms of its potential impact on pest and disease risk due to dispersal, using an index summarizing key metrics for cropland connectivity risk based on a gravity model, with associated uncertainty quantification, and iii) use the network structure to identify geographic priorities for surveillance and management of emerging pests and diseases, and for saturation of endemic species. We discuss the geographic spread of diseases in the context of cropland connectivity using as an example some key diseases and pests, including banana bunchy top disease, Xanthomonas wilt of bananas, potato yellow vein, and Guatemalan potato tuber moth.

## Evaluating the potential role of locations in spread networks

### Network metrics to evaluate invasion and saturation risk

We consider a set of network metrics that have often proven useful for evaluating the role of nodes in network processes. To simplify comparisons, we also summarize across metrics in a cropland connectivity risk index (CCRI). To emphasize the importance of the node as a bridge, the index emphasizes betweenness centrality, based on the number of shortest paths through the network that include the node being evaluated. The other half of the weight is given to other metrics that measure how well connected a node and its neighbors are: node strength (the sum of a node’s link weights), the sum of a node’s nearest neighbors’ node degrees (sum of the number of links associated with each nearest neighbor), and Eigenvector centrality (giving each node a score proportional to the sum of the scores of its nearest neighbors and more distant neighbors). The summary index (the CCRI) was calculated as a weighted sum of 1/2 betweenness centrality, 1/6 node strength, 1/6 sum of nearest neighbors’ node degrees, and 1/6 Eigenvector centrality, where each of the four metrics was scaled before summing by dividing by the maximum value observed for that metric. The weighting emphasizes betweenness because betweenness will particularly capture a potential role as a bridge that is not obvious when individual cropland area is considered alone, and also to include connectedness of a node at different scales.

### Illustration of features captured by the cropland connectivity risk index

Before we consider cropland connectivity risk for individual global crop systems, here is an illustration of how the four metrics described above (betweenness, node strength, sum of nearest neighbors’ node degree, and eigenvector centrality) capture different elements of cropland connectivity. Suppose this hypothetical map represents cropland density for a target crop species (Fig. 3A). In this example, most of the cropland units have a low crop proportion (indicated by yellow shading) while one unit has a high crop proportion (indicated by blue shading). A network (Fig. 3B) is constructed using the gravity model described above, from the corresponding data for the cropland map (Fig. 3A), based on one parameter combination. (More details about the exponential and inverse power law models are given below.) The high crop-density location on the map is represented by the blue node in the network while the other nodes represent the other land units where the crop was present. Note that this is the network for one particular parameter combination, while the later components of Fig. 3 represent summaries across uncertainty quantification, as described more below. The sum of the node degrees for nearest neighbors (Fig. 3C) captures how well connected the nodes’ neighbors are. Node strength (Fig. 3D) indicates a node’s importance in terms of how connected it is to its neighbors. Betweenness centrality identifies nodes acting as bridges to connect other regions in the network (Fig. 3E). Eigenvector centrality (Fig. 3F) shows how well connected a node is through nearest neighbors, their neighbors, and beyond. The cropland connectivity risk index (CCRI; Fig. 3G) is the weighted mean of these metrics. In addition to the high-risk locations with high crop density, other locations with high risk because of their role as bridges were identified. The results of an uncertainty quantification for the CCRI in this hypothetical map are also shown (Fig. 3H, 2J-L), illustrating how a summary across parameter combinations (beyond the combination illustrated in Fig. 3B.) can reveal other features of a cropland landscape. Finally, we identify locations where CCRI rank among locations is higher than the rank based solely on crop density (Fig. 3I). These are locations where the network analysis reveals potentially important roles for a location that would not be apparent in a simpler point-wise analysis of crop density.

**Figure 3.**
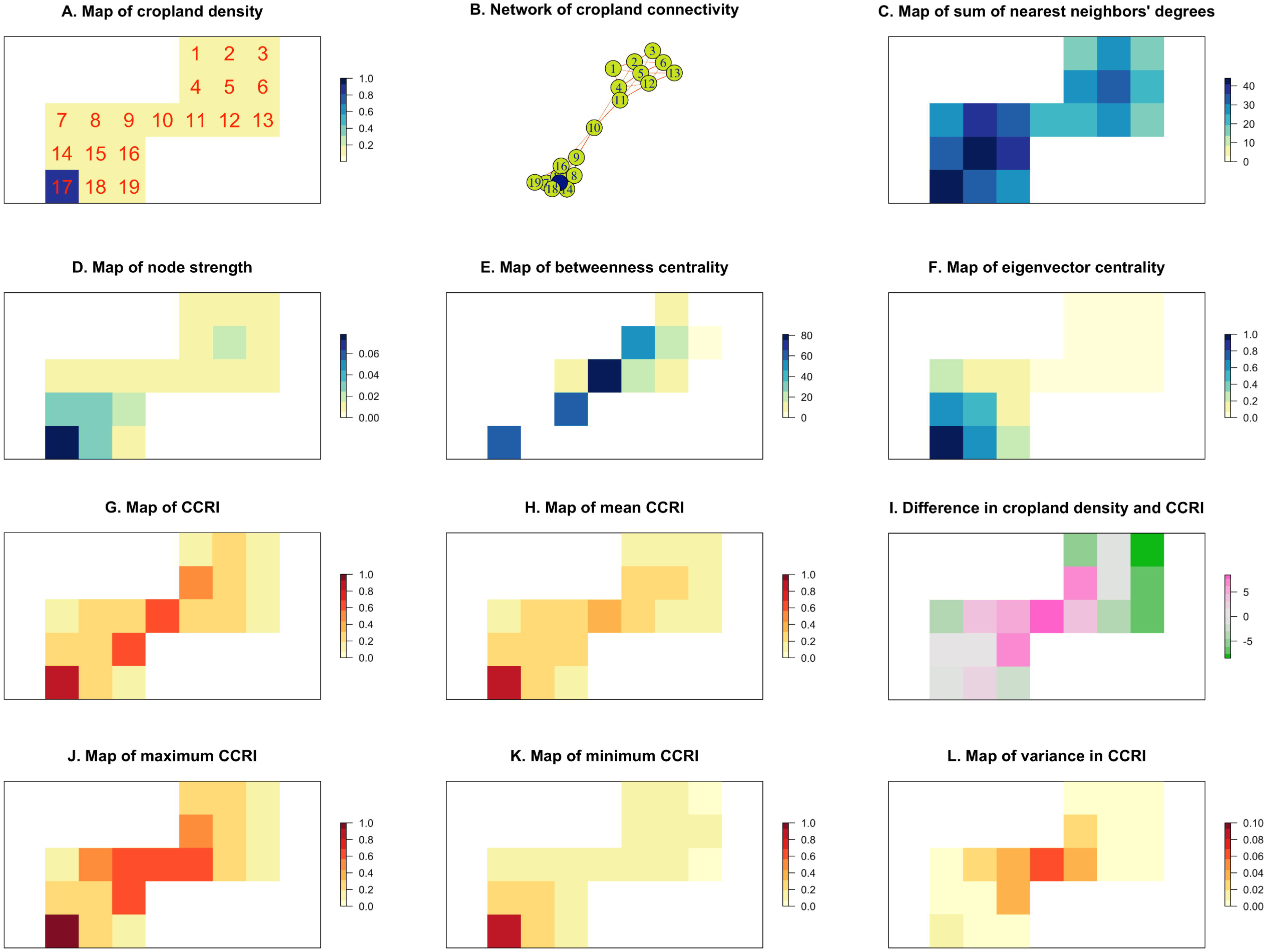
An illustration of the evaluation of cropland connectivity risk for a simple hypothetical scenario. A. The map of cropland density indicates the fraction harvested area for a crop species in a hypothetical small region, where white areas have none of the crop species, green areas (1-16 and 18-19) have a low fraction of land planted to the crop species, and the blue area (17) has a high fraction planted to the crop species. B. The network of cropland connectivity that corresponds to the map in A, indicating the links for one set of threshold parameters (negative exponential function with *γ* = 0.7 was used to calculate the link weight, and a threshold of 0.001 was used to determine whether a link exists). The high density region (blue node 17) and bridging region (green node 10) are indicated. C. A map of the sum of nearest neighbors’ degrees for the network in B. Nearest neighbors are those with direct links to a reference node, and node degree is the number of links to that node. D. A map of node strength for the network in B. Node strength is the sum of the weights of links to a reference node. E. A map of betweenness centrality for the network in B. Betweenness centrality indicates the number of shortest paths in the network that pass through a reference node. F. Eigenvector centrality for the network in B. Eigenvector centrality is a measure of how well connected a node is in terms of immediate neighbors, their neighbors, etc. G. The cropland connectivity risk index (CCRI) for the network in B. CCRI is a weighted mean of the four measures of connectedness in maps C through F. Note that this is the CCRI for one parameter combination, while additional combinations are illustrated in the supplemental material. H. A map of the mean CCRI from uncertainty quantification for networks corresponding to map A for a range of parameter combinations (Table1). I. The difference between ranked values of the mean CCRI from a sensitivity analysis, and the ranked values in map A. This difference indicates locations where the fraction of land planted to the crop species does not capture all the features of connectivity in the CCRI. Pink indicates regions where the CCRI is higher than the ranked values of map A, and green indicates regions where the CCRI is lower. J. Map of the maximum CCRI from the sensitivity analysis. K. Map of the minimum CCRI from the sensitivity analysis. L. Map of the variance in CCRI across realizations in uncertainty quantification.

### Global cropland area

Now we expand this cropland connectivity analysis to global cropland. We analyzed two standard data sets representing the global geographic distribution of individual crop species: data representing conditions circa 2000 from Monfreda et al. (2008), referred to here as the “Monfreda” data set, and IFPRI’s Spatial Production Allocation Model (SPAM) data 2005 v3.2 (IFPRI and IIASA 2016), referred to here as the “SPAM” data set. Each of the two global cropland datasets was analyzed individually using the methods described below. We also evaluated a combined dataset, “Monfreda & SPAM”, containing the mean of the harvested area fraction from the two data sources.

For each of these three datasets, we evaluated the harvested cropland area for banana, cassava, potato, sweetpotato, and yam. Here we use the term “banana” to refer to both bananas and plantains (Beed et al. 2012). Data were spatially aggregated by finding the mean harvested area for each crop across 24 × 24 grids of the original 5 min x 5 min grids, for a resolution of 120 min x 120 min (2°x 2°). We used two methods to calculate the mean of the crop harvested area per grid: the “land mean” (the sum of the harvested area fractions within an aggregated 2°x 2°unit divided by the number of 5 min x 5 min units within the aggregated unit that contain only land) and the “total mean” (the same sum divided by the total number of 5 min x 5 min units within an aggregated 2°x 2°unit). These two formulations of the mean are different primarily on coastlines and for islands; the uncertainty quantification below addresses both formulations. To focus on more important production areas, we considered three threshold values for inclusion of nodes in the analysis: 0.0015, 0.002, and 0.0025 mean proportion cropland harvested area. We described the risk for pathogen and pest movement between each pair of nodes as a function of the distance between the nodes and the cropping density associated with the nodes.

### Model of risk of movement between geographic nodes

We constructed adjacency matrices where the entry for each pair of nodes was a function of the distance between the two nodes and the cropping density at the two nodes. The distance effect on the risk was calculated as a function of the Vincenty ellipsoid (Hijmans et al. 2017c) distance between nodes *i* and *j* (*d*_*ij*_) as either an inverse power law function *d*_*ij*_^*-β*^, or a negative exponential function exp(^*-γd*^_*ij*_) (Campbell and Madden 1990, Gregory 1968, Madden et al. 2007, Mundt et al. 1999, Mundt et al. 2009b, Severns et al. 2014). For convenience, the distance in meters was further scaled by dividing by 111,319.5 (one degree Vincenty ellipsoid distance in meters at the equator). Higher values of the parameters *β* and γ reflect lower likelihood of long-distance dispersal. Three levels (0.5, 1.0, and 1.5) of parameter *β* were considered for the inverse power law function in uncertainty quantification (Table 1), generating a 31.6%, 10.0%, and 3.16% chance of movement, respectively, across a distance of 10 km (a 5-min resolution cell is approximately 10 by 10 km at the equator). Five levels (0.05, 0.1, 0.2, 0.3, and 1.0) of parameter γ were evaluated in uncertainty quantification, generating corresponding chances of movement across a distance of 10 km of 60.6%, 36.8%, 13.5%, 4.98%, and 0.0045%, respectively, before adjustment for the proportion harvested area in the two linked pixels. The risk due to greater cropland area for any two nodes *i* and *j* was accounted for using a gravity model (Jongejans et al. 2014), by multiplying together the mean cropland area (*c*) associated with each of the nodes (*c*_*i*_ *c*_*j*_). Thus, in the first step the weights in the adjacency matrix indicating the overall risk of movement between two geographic nodes were *c*_*i*_*c*_*j*_*d*_*ij*_^*-β*^ for the inverse power law function and *c*_*i*_*c*_*j*_*exp* ^*-γd*^ _*ij*_ for the negative exponential function. In the uncertainty quantification we also evaluated results across three threshold minimum values (0.001, 0.0001, and 0.00001) for entries in the matrix individually and set weights below that to be zero. Network models and metrics for the cropland connectivity for each of the five crops were analyzed for the Eastern and Western Hemispheres separately. We used the igraph package (Csárdi and Nepusz 2006) in the R programming environment (R Core Team 2018) to evaluate the networks, and the geographic data analysis and modeling package raster (Hijmans et al. 2017b). Other R packages used were rgdal (Bivand et al. 2018a), dismo (Hijmans et al. 2017a), expm (Goulet et al. 2017), maptools (Bivand et al. 2017), rrcov (Todorov 2018), rworldmap (South 2016), mapdata (Becker et al. 2018a), sp (Pebesma et al. 2018), maps (Beckeret al. 2018b), rgeos (Bivand et al. 2018b), geosphere (Hijmans et al. 2017c), viridis (Garnier et al. 2018), and RColorBrewer (Neuwirth 2014).

**Table 1.**
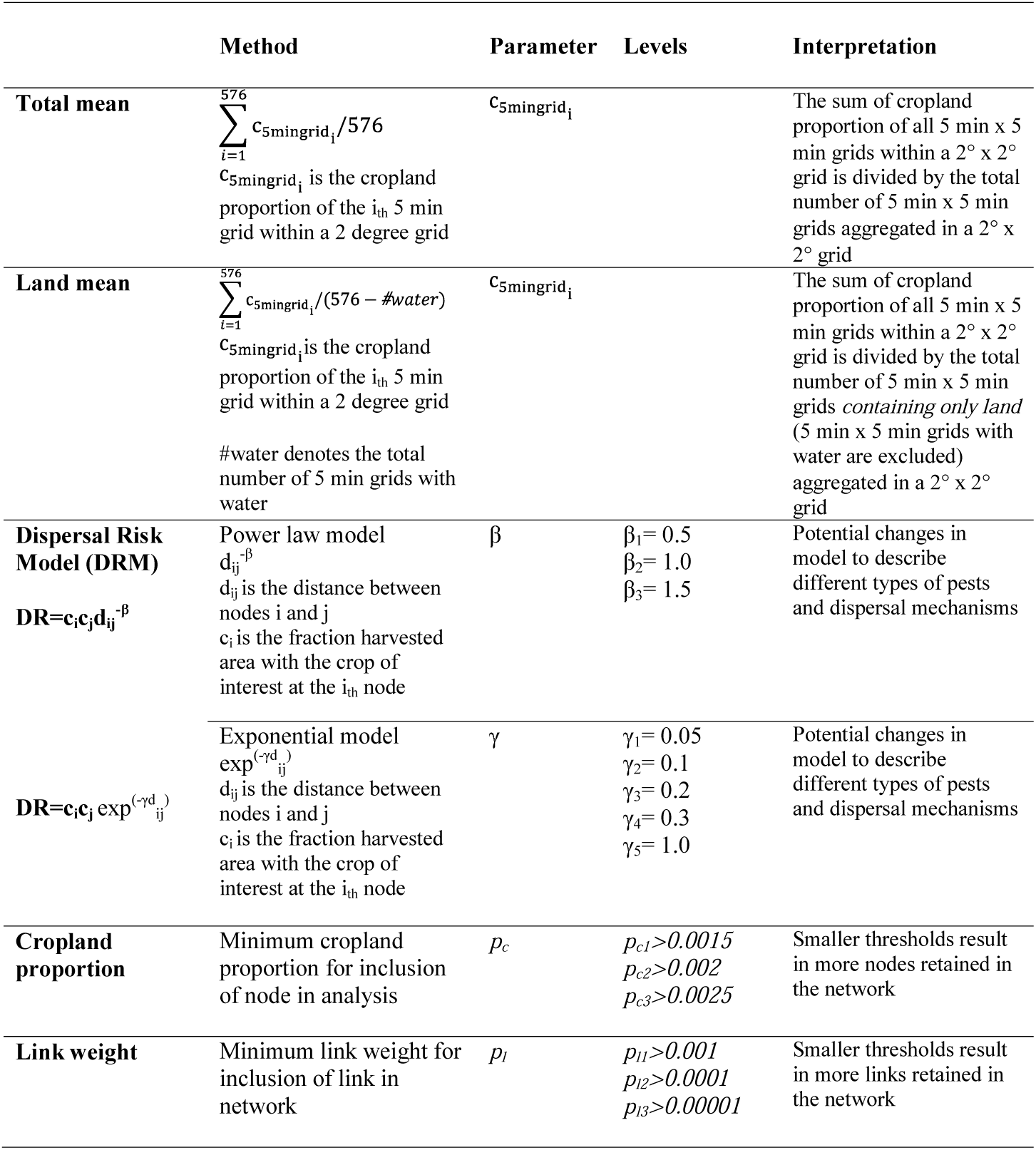
Components of a summary measure of cropland connectivity risk index (CCRI) that were evaluated in uncertainty quantification. Each combination of the levels of the values indicated was evaluated. The combinations included varying the form of mean (total mean or land mean) and varying the dispersal model (inverse power law or negative exponential), as well as the parameters of the model selected.

### Uncertainty quantification and data quality

We performed an analysis of model sensitivity to parameter shifts, to i) evaluate how consistent results were under changes in model parameters, and ii) determine which nodes had high cropland connectivity risk across all or most model scenarios, and which nodes had high risk only for a limited range of model scenarios. Based on the combinations of functions, thresholds, and parameters used, 144 cropland connectivity risk index maps were generated for each crop (Table 1). For each cell in the maps, we calculated the mean, max, min, and variance across the 144 maps. And, for reference, we summarize the data quality assessment provided by Monfreda et al. (2008; Fig. S3). We also compared how locations rank based on CCRI to how they rank based on harvested crop fraction (crop density). Those locations where the CCRI rank is substantially higher than the crop density rank are locations that might have particularly important roles in epidemic spread but that would not be identified if analysis looked solely at crop density.

### Interpreting maps of cropland connectivity risk

Mapped areas with a higher CCRI are likely to have higher risk for dispersal of pathogens or pests of banana and plantain, cassava, potato, sweetpotato, and yam, based on cropland connectivity. Cropland connectivity is a risk factor for movement through wind dispersal, active pest movement, vector movement, seed exchange, farm tools, or trade. The locations we identified are candidates for prioritizing surveillance and mitigation programs (Smolinski et al. 2003, Woolhouse et al. 2005), especially if information about weather conduciveness to invasion, e.g., suitable temperature (Kroschel et al. 2016), and other risk factors such as documented trade patterns (Andersen et al. 2019), also support the high risk designation.

A cropland connectivity risk index will often be an important component of integrated geographic risk assessment, along with weather/climate risk factors, genetic resistance deployment, and trade. We demonstrated how a cropland connectivity risk index can be designed to go beyond simply identifying as high risk those land units which have high crop fraction, especially if the index captures how locations may function as bridging by incorporating a measure such as betweenness centrality. Land units with high crop fraction <u>will</u> tend to be identified as high risk, although less so to the degree that they are isolated, because the gravity model weights crop fraction in evaluation of the probability of movement. However, the cropland connectivity risk index also identifies locations that have an important role as bridges between cropping regions (pink regions in Figs 3I and 5), even if the cropping density within the bridge land units is not particularly high. Thus, analysis of cropland connectivity can identify additional risk areas based on the larger landscape, beyond those identified through a simple unit-by-unit scan for high cropping density.

We present analyses for the crop species individually. However, as an overall measure of cropland connectivity risk for pathogen and pest invasion and saturation for these crops, a cross-crop index constructed by adding the individual species risk indices may be useful for evaluating strategies for general purpose surveillance and management strategies for the set of crops. Combining across host species might also be useful for special cases where a target pathogen or pest uses multiple host species.

## Cropland connectivity and the risk of major pests and diseases of roots, tubers, and bananas

### Banana and plantain

A combination of high mean CCRI and low variance in CCRI in uncertainty quantification was observed for central, north central and southern Uganda, northwest Tanzania, Rwanda, Burundi, the Inter-Andean valleys in Colombia, central and western Ecuador, and Haiti. The highest global CCRI was found in the border region of Uganda, Rwanda, and Tanzania (Fig. 4A). The combination of high mean and low variance indicates that these locations are more likely to have a high risk across all model assumptions evaluated. The CCRI rank was substantially higher than the rank based on cropland density alone in multiple locations in Africa, particularly in Tanzania (Fig. 5A).

**Figure 4.**
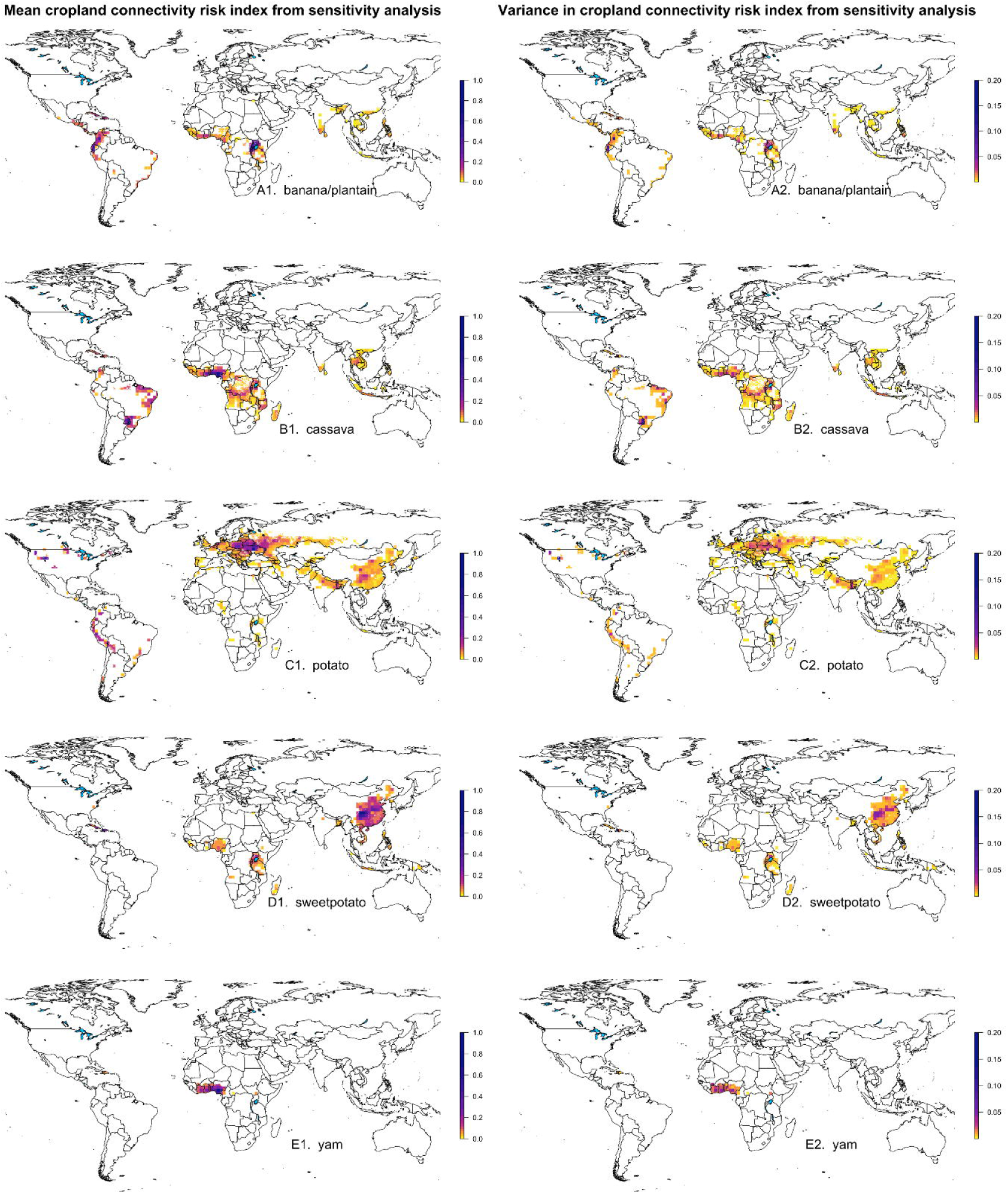
The mean and variance observed for cropland connectivity risk index (CCRI) components in uncertainty quantification for banana/plantain, cassava, potato, sweetpotato, and yam (based on the mean of crop density estimates from Monfreda et al. (2008) and SPAM (You et al. 2017)).

**Figure 5.**
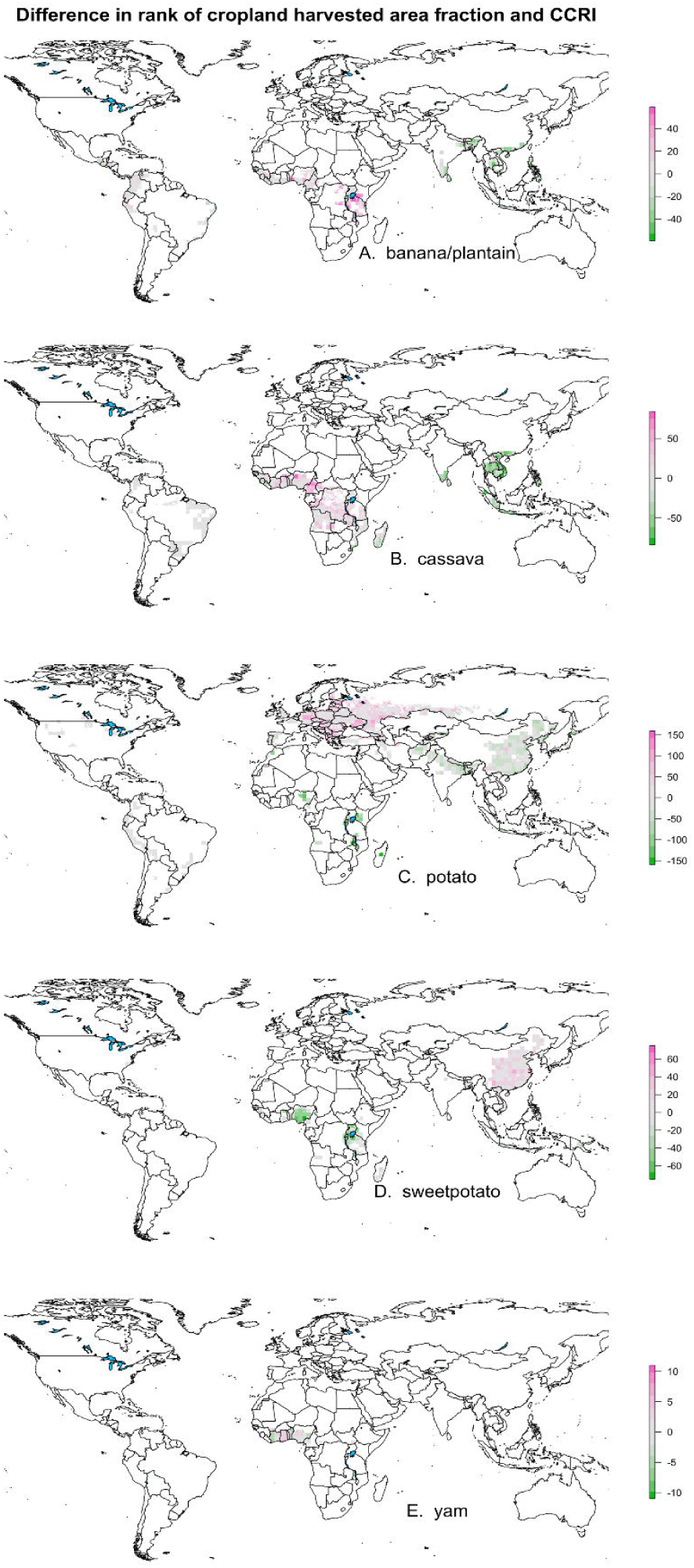
Maps of the difference in cell rank between harvested area fraction and the mean cropland connectivity risk index (CCRI) for banana/plantain, cassava, potato, sweetpotato, and yam (based on the mean of crop density estimates from Monfreda et al. (2008) and SPAM (You et al. 2017)). Locations where the CCRI has substantially higher rank than the crop density are locations that could have important roles in spread that would not be detected if analyses were limited to crop density.

Two banana diseases illustrate how pathogen invasion and spread can be linked to cropland connectivity. Banana bunchy top disease, caused by Banana bunchy top virus (BBTV, genus *Babuvirus*), causes devastating losses. Phylogenetic studies of BBTV spread in Africa, along with farmers’ observations, suggest dual introduction events for BBTV – in Egypt and then in the Democratic Republic of Congo (DRC) – before further virus spread in Sub-Saharan Africa (Kumar et al. 2011, Leung et al. 2015). Movement of planting material, and long-distance spread facilitated by migrant workers, likely have contributed to the gradual expansion of BBTV in SSA. Ubiquitous distribution of the vector, the banana aphid (*Pentalonia nigronervosa*), contributed to local spread of the virus. Xanthomonas wilt of banana (caused by *Xanthomonas campestris* pv. *musacearum*) often causes yield reductions of up to 100% (Ndungo et al. 2006, Tripathi et al. 2009, Tushemereirwe et al. 2004). This disease is mainly spread by infected planting material, insect vectors, farm tools, browsing animals, and occasionally by bats, birds, and weevils (Gold and Bandyopadhyay 2005, Tinzaara et al. 2006, Were et al. 2015, Yirgou and Bradbury 1974). The disease had been limited to the Ethiopian enset growing belt until 2001 when it appeared in banana fields in central Uganda and eastern DR Congo (Ndungo et al. 2006, Tushemereirwe et al. 2004). Highly connected and susceptible Musa ABB type production systems dominate in lower elevation central Uganda, where there is high insect activity, and this combined with often poor plot management levels likely produced the particularly fast spread of the disease. In contrast, and despite a high level of connectivity of banana cultivation zones, disease spread has been slower in the Albertine rift valley mountainous region of east DR Congo. This is probably mainly due to elevation effects on insect vector transmission and the predominance of AAA-EA highland banana types, which are less susceptible to insect vector transmission. In this region, disease spread mainly occurs through garden tool use. The adjacent Congo basin lowlands to the west of the rift valley are dominated by dense tropical forests, with banana/plantain only in villages along road or river axes, and often low connectivity (G. Blomme, personal observation). As a result, disease spread into the Congo basin has, over a period of nearly 20 years, been very limited.

### Cassava

The CCRI for cassava was particularly high throughout southern Nigeria, south central Ghana, western Burundi, northern Brazil (Bahia and Amazonas states), southern Brazil (Parana and Mato Grosso states), southwest Paraguay, and northeast Argentina (Fig. 4B). The CCRI rank was substantially higher than the rank based on cropland density alone in locations in Nigeria, Cameroon, and the Democratic Republic of Congo (Fig. 5B).

Cassava frogskin disease is an economically important disease of cassava across Latin America and the Caribbean, where up to 90% yield loss was reported in severely infected fields in the 1980s (Pineda et al. 1983). The disease has been associated with several pathogens, and transmission is typically via asymptomatically infected cassava planting material (Alvarez et al. 2009, Carvajal-Yepes et al. 2014). Until 1971 the disease had been found only in Colombia, but since then it has been reported in Panama, Costa Rica, Venezuela, Peru and Brazil, (Calvert et al. 2012, Calvert and Thresh 2002, Chaparro-Martinez and Trujillo-Pinto 2001, Di Feo et al. 2015). A more recent threat to cassava production is the re-emergent ‘cassava common mosaic disease’ (CCMD), caused by the mechanically transmitted potexvirus Cassava common mosaic virus. Originally reported to cause significant yield losses in southern Brazil since the 1940s (reviewed in Lozano et al. (2017)), recent outbreaks of CCMD have been reported in Peru and Argentina (Di Feo et al. 2015, Fernandez et al. 2017, Zanini et al. 2018). Because of the likely spread of these pathogens through asymptomatic planting material, it is probable that patterns of spread throughout Latin America are related to high intensity areas of production. For example, CCMD is reported in high intensity areas of production in northeast Peru and northeast Argentina, where high CCRI was observed in our analysis.

In contrast to CMD (discussed earlier), cassava brown streak disease (CBSD) is less readily spread by vectors. Its rate of spread into Central Africa was likely also slowed due to lack of cassava cropland connectivity, associated with the massive barrier of the great forests of the Congo Basin. High levels of cassava cropland connectivity in West Africa, from western Cameroon westwards, suggest that if CBSD is introduced to this area, there is likely to be rapid spread further westwards. Both cassava pandemics are being driven by super-abundant populations of the whitefly vector, *Bemisia tabaci*. Cropland connectivity is also likely to be important for Bemisia whitefly populations, because the whitefly genotypes occurring on cassava have a strong preference for this crop (Wosula et al. 2017).

### Potato

The CCRI for potato was particularly high in northcentral Europe, including northern and central Ukraine, central Poland, central and southern Belarus, and southwestern Russia. CCRI was also high in locations in the United States (Idaho, Washington, Colorado, and the northern Great Lakes region), New Brunswick in Canada, locations in Peru and central Colombia, and central China (Fig. 4C). In Africa, the Lake Kivu region has a high CCRI. The CCRI rank was substantially higher than the rank based on cropland density alone in multiple locations in Eastern Europe and Eastern China (Fig. 5C).

Potato yellow vein is an important potato disease in the northern Andean region caused by Potato yellow vein virus (PYVV). PYVV is transmitted by the greenhouse whitefly (*Trialeurodes vaporariorum*), through seed potato and underground stem-grafts (Salazar et al. 2000). Originally reported in northern Ecuador and west central Colombia, the virus has spread, probably via infected seed tubers, throughout the central Andes, particularly to the most important potato-producing areas of Colombia (Guzmán et al. 2006), Venezuela, and northern Peru. Interestingly, PPVV has not moved further south over the last 20 years despite predicted favorable conditions for whiteflies in these regions (Gamarra et al. 2016). Likely a gap with reduced cropland connectivity and limited potato seed exchange between northern and southern Peru has contributed to this lack of spread, although other factors, such as cultivar resistance, may also be important factors. Central Peru should be a priority area for monitoring to prevent further spread of the disease to southern Peru and Bolivia, whereas spread to other regions is likely only possible through long distance transport of infected potato.

The potato tuber moth, *Tecia solanivora*, is a challenging potato pest in Central and South America (Kroschel and Schaub 2013). Guatemala is understood to be the country of origin. In 1970, the pest was accidently introduced with infested seed into potato growing regions of Costa Rica; in 1983, into Venezuela, and, in 1985, into Colombia. In 2010, *T. solanivora* was reported for the first time from southern Mexico, and in 1996 from Ecuador. In 1999, *T. solanivora* appeared on Tenerife, Canary Islands. Since then the pest has been considered a major threat to potato crops throughout southern Europe, and was listed as a quarantine pest by the European and Mediterranean Plant Protection Organization (EPPO 2005). Schaub et al. (2016) confirmed the suitable climatic conditions in southern Europe. However, as *T. solanivora* is strongly monophagous and potato is its only host plant, the movement of infested seed is the main potential pathway of its spread into new potato growing regions, especially if there is not a high level of cropland connectivity among the regions. The pest was therefore contained in Tenerife for many years, before it was first detected in mainland Spain in Galicia in 2015, and in Asturias in 2016 (Jeger et al. 2018). In contrast, the South American tomato leafminer, *Tuta absoluta*, after its transatlantic invasion and first detection in Spain in 2006, rapidly spread across southern Europe, Africa, and Asia (https://gd.eppo.int/taxon/GNORAB/distribution). Compared to *T. solanivora*, also a Lepidopteran pest of potato, the very rapid spread of *T. absoluta* is likely due at least in part to its very wide host range in the Solanaceae, and the very high level of connectivity of these combined species. Hence, considering the low level of connectivity of the potato crop in many regions, e.g, in southern Europe and much of Africa, it will likely be more difficult for a monophagous potato pest such as *T. solanivora* to invade new potato growing regions.

### Sweetpotato

The CCRI for sweetpotato was high in locations in central China, the Caribbean (Haiti and the Dominican Republic) (Fig. 4D), and in central Uganda, with central China having the highest ranked global risk. The CCRI rank was substantially higher than the rank based on cropland density alone in multiple locations in China (Fig. 5D).

In sweetpotato, several weevils important to yield loss exist worldwide. Sweetpotato viruses such as Sweet potato chlorotic stunt virus (SPCSV), Sweet potato feathery mottle virus and some begomoviruses are already present globally, whereas other viruses such as Sweet potato mild mottle virus are found only in certain regions. Some strains of sweetpotato viruses, such as the severe EA strain of SPCSV, are geographically localized. Movement of planting material (sweetpotato vines) through trade can cover long distances (Rachkara et al. 2017). Whereas some sweetpotato pests such as viruses are easily spread through planting material (Gibson and Kreuze 2015) and can form permanent reservoirs in wild host species (Tugume et al. 2008, Tugume et al. 2013, Tugume et al. 2016), others, such as weevils, are not readily spread through planting material, have no known alternative hosts, and are unable to travel long distances by themselves.

### Yam

Of the crops evaluated, yam had the lowest overall global harvested area. The highest CCRI observed for yam was in locations in southcentral Nigeria, Benin, Togo, Ghana, and the Ivory Coast, along with locations in the Caribbean including Haiti (Fig. 4E). The CCRI rank was substantially higher than the rank based on cropland density alone in locations in eastern Nigeria, Togo, western Ivory Coast, and the Dominican Republic (Fig. 5E).

Yam is a multispecies crop grown for its tubers by millions of smallholder farmers in West Africa. Nearly 94% of global edible yam production is in West Africa (Benin, Cameroon, Côte d’Ivoire, Ghana, Nigeria and Togo), and Nigeria alone produces 66% of global production (FAOstat 2016). Major constraints to yam production in West Africa are mosaic disease caused by Yam mosaic virus and Yam mild mosaic virus (genus Potyvirus), and anthracnose caused by *Colletotrichum gloeosporioides*. Damage to yam by nematodes – *Scutellonema bradys, Pratylenchus* spp., and *Meloidogyne* spp. – is responsible for significant pre- and postharvest deterioration of tubers. All these agents are endemic in all the yam production regions in West Africa, so saturation is the main consideration *within* that region, and anecdotal evidence clearly links the spread of yam viruses and nematodes to the movement of planting material in West Africa.

### Incorporating cropland connectivity in risk assessments

These analyses illustrate general cropland connectivity risk across a large spatial extent and for a fairly coarse spatial resolution. Follow-up analyses for specific locations and particular pathogen or pest species may be useful, when more detailed data are available for mapping cropland fraction and for selecting appropriate functions to describe dispersal kernels for specific time scales, and potentially other factors. The results presented here have a greater confidence for certain crops such as potato, and certain regions, based on the quality and quantity of the original data available for assembly by Monfreda et al. (2008) and in SPAM 2005 v3.2 (IFPRI and IIASA 2016). Examples of the application of network analysis to invasions of particular species include *Phytophthora ramorum* (Harwood et al. 2009, Shaw and Pautasso 2014) and *Phakopsora pachyrhizi* (Sanatkar et al. 2015, Sutrave et al. 2012). The role of a land unit will depend on the species of pathogen or pest being considered, and its dispersal kernel. A particular land unit evaluated for species that tend to move only short distances might be isolated, while for species that tend to move longer distances it might be an important bridge node (Calabrese and Fagan 2004). Individual pathogen or pest data and implications for trade can be disseminated via actively updated Regional Pest Risk Assessment working documents, to support standards established by the International Plant Protection Convention to prevent introduction, establishment and spread of pests and diseases, implemented by National and Regional Plant Protection Organizations (Beed et al. 2013, IPPC 2012, Miller et al. 2009).

Other useful points for future research to refine cropland connectivity risk assessments include the following. The current analysis is based on geographic data for 2000 (Monfreda et al. 2008) and 2005 (IFPRI and IIASA 2016), so areas where crop densities have increased rapidly in recent years, are not represented in these global maps yet. An important example is cassava production in SE Asia, where production has quickly expanded and is now experiencing an invasion of cassava mosaic disease (Delaquis et al. 2018, Wang et al. 2015). The cropland density data are summarized across global data sets that vary widely in quality from region to region. The resolution we selected for our analyses was intended to represent a compromise – avoiding too high a spatial resolution because it might have little data to back it up in many regions, and also avoiding too coarse a resolution that might obscure the roles of specific regions. Where more complete data are available or can be collected, more detailed and higher resolution analyses can be performed. Likewise, the current analysis does not take into account geographic features that could have important effects on the likelihood of active or passive movement of pathogens and pests, or weather features such as wind patterns (Sutrave et al. 2012). Roads and rivers may increase pathogen movement, while other water bodies, deserts, and mountains may isolate nodes (Meentemeyer et al. 2012). And the distribution of individual crop species captures only some aspects of risk for many pathogens and pests that can use multiple host species. Conversely, if resistance genes are widely deployed, pathogens and pests may only be able to use a subset of the planted fraction (Brown and Hovmoller 2002, Garrett et al. 2017). Extreme weather patterns may be responsible for many important regional or global introductions of pathogens and pests, such as the potential introduction of soybean rust to the US in hurricane Ivan (Schneider et al. 2005). Flooding may move some soilborne pathogens to new locations. Finally, heterogeneity in time may alter patterns of cropland connectivity. Markets may drive longer term trends in planting patterns, and for shorter season crops such as potato and sweetpotato, geographic heterogeneity in planting seasons may disrupt the cropland connectivity suggested when seasons are aggregated.

Two regions in Africa, the Great Lakes Region and the region between Ghana and Nigeria, have high cropland connectivity risk for multiple crops. East African cassava mosaic virus-Uganda (EACMV-Ug) emerged in Uganda and caused famine in East, Central and West Africa (Anderson et al. 2004), and wheat stem rust race Ug-99 also emerged in Uganda (Pretorius et al. 2000). It is an interesting open question if the geographic position of Uganda in cropland networks had some influence on disease emergence, or if it was simply a matter of higher sampling effort that made detection more likely.

Areas identified with low connectivity and low risk can also be prioritized to produce disease-free seed. This has often happened naturally, for example where particularly dry regions that are not optimal for crop production, and thus are often isolated from other crop production regions, may produce seed with low disease risk. Seed production areas play an outsize role in the risk of disease spread, even where cropping density is low. The risk of pathogen spread through seed networks is a key component for integration with risk based on cropland connectivity (Andersen et al. 2019, Garrett et al. 2018). The movement of pathogens through the international seed trade is an important risk factor for many crops (Anderson et al. 2004, Rodoni 2009, Wylie et al. 2008). In Sub-Saharan Africa, movement of plant material and farming tools is a key factor for the dispersal of banana diseases such BXW (Beed 2014, Tripathi et al. 2009) and through cuttings for cassava virus diseases, particularly CBSD (Bock 1994, Legg et al. 2015). The US late blight pandemic in 2009 was caused by the movement of infected tomato plants via trade from a single national supplier (Fry et al. 2013). During 2009-2010, an epidemic of *P. infestans* in tomato was reported in southwest India with the suggestion that the pathogen was introduced via seed potato imported from the UK and Europe before 2009 (Chowdappa et al. 2013). Cropland connectivity is likely to capture at least a portion of the risk associated with movement of seed, transplant, and agricultural equipment, to the extent that trade and movement of equipment and agricultural workers tends to follow a path through areas that produce a particular crop. Of course, at the same time that cropland connectivity represents a risk for the spread of pathogens and pests, connectivity may also sometimes confer advantages for efficiency in deployment of equipment and personnel, as well as marketing of seed and produce.

In summary, cropland connectivity constitutes a risk component important for most pests and diseases. It can complement risk assessments based on the effects of climate, genetic resistance, and formal trade networks. The integration of these risk assessment layers will make the best use of available data to evaluate risk and guide targeted surveillance and mitigation in global strategies (Carvajal-Yepes et al. 2019). Uncertainty quantification can help in interpreting analyses when information about dispersal kernels is not available, or when a more general analysis is desired, and in targeting data collection to address the most important data for improving key parameters. Availability of high quality data related to cropping density, deployment of genetic resistance, and weather patterns will improve risk assessments. Developing optimal methods for integration across cropland connectivity and other risk data layers is an important challenge for future risk analysis, and offers the promise of more effective risk assessment in the future.

## Acknowledgements

This research was undertaken as part of the project “Management of RTB-critical pests and diseases under changing climates, through risk assessment, surveillance and modeling”, funded by the CGIAR Research Program on Roots, Tubers and Bananas (RTB) and supported by CGIAR Fund Donors https://www.cgiar.org/funders/, Bill and Melinda Gates Foundation grant OPP1080975, the CGIAR Research Program on Climate Change and Food Security (CCAFS), USDA APHIS grant 11-8453-1483-CA, US NSF Grant EF-0525712 as part of the joint NSF-NIH Ecology of Infectious Disease program, US NSF Grant DEB-0516046, and the University of Florida. We appreciate a helpful discussion with N. Ramankutty.

